# Evolutionary trajectories determine the feasibility of collateral sensitivity based antibiotic treatment strategies in critical bacterial pathogens

**DOI:** 10.1101/2025.06.03.657621

**Authors:** Vatsala Chauhan, Lisa Enkvist, Yuliia Chukhareva, Carl Damell, Eric Cruz Davila, Tasnim M. Islam, Greta Melander, Ellen Paulsson, Adam Sundell, Emily Zweifel, Anna Klercker, Anna Knöppel, Gerrit Brandis

**Author notes:** Corresponding author: Gerrit Brandis.

## Abstract

The rise of antibiotic resistance among pathogenic bacteria necessitates innovative therapeutic strategies. A promising technique is the use of collateral sensitivity where resistance to one antibiotic increases susceptibility to another. In this study, we explored the clinical relevance of collateral sensitivity through experimental evolution and genetic engineering in six critical bacterial pathogens using 23 distinct antibiotics. Our in-depth analysis of *Escherichia coli* showed that clinically relevant resistance mutations did not confer collateral sensitivity to the tested antibiotics. We were able to identify at least three new classes of ciprofloxacin-resistance mutations that cause collateral sensitivity to multiple antibiotics. However, these mutations incur significant fitness costs and are absent in ciprofloxacin-resistant clinical isolates. Our further analysis showed that the development of collateral effects differed significantly between the tested species. Most species showed development of collateral sensitivity to gentamicin during ciprofloxacin-resistance evolution but *Acinetobacter baumanii* developed collateral resistance instead. Overall, *Pseudomonas aeruginosa* showed the most consistent development of collateral sensitivity among the tested species, highlighting it as a promising candidate for the use of collateral-sensitivity-based treatment strategies. Our findings provide insights into the potential of collateral sensitivity as a therapeutic strategy and contribute to the development of more effective antibiotic treatment regimens.

## Introduction

The rise of antibiotic resistance among pathogenic bacteria poses a significant threat to global public health, with current forecasts estimating over 10 million deaths per year attributable to or associated with antibiotic resistance by 2050 (GBD 2021 Antimicrobial Resistance Collaborators 2024). This alarming trend necessitates the establishment of novel therapeutic strategies to combat drug-resistant infections. One promising approach is the exploitation of collateral sensitivity, an evolutionary tradeoff in which resistance to one antibiotic increases susceptibility to another (Roemhild and Andersson 2021; Maltas, et al. 2025). This phenomenon offers a potential strategy for designing treatment regimens to limit the emergence and spread of antibiotic resistance (Mahmud and Wakeman 2024).

Collateral sensitivity has been observed in various prokaryotic and eukaryotic species in laboratory evolution experiments. These include Gram-negative bacteria (Barbosa, et al. 2017; Imamovic, et al. 2018; Podnecky, et al. 2018; Hasan, et al. 2023), Gram-positive bacteria (Gonzales, et al. 2015), *Mycobacterium tuberculosis* (Trigos, et al. 2021), and *Candida auris* (Carolus, et al. 2024). Molecular mechanisms underlying collateral sensitivity have been identified in some cases. For example, in *Escherichia coli*, tigecycline resistance causes collateral sensitivity to nitrofurantoin due to increased expression of nitroreductase enzymes, increased drug uptake rates, and increased drug toxicity (Roemhild, et al. 2020). In *Pseudomonas aeruginosa*, ciprofloxacin resistance caused by overexpression of the MexCD-OprJ eflux pump results in major changes in cell envelope physiology, leading to collateral sensitivity to ß-lactam antibiotics (Mulet, et al. 2011). These findings suggest that collateral sensitivity can be leveraged to develop antibiotic cycling or combination therapies that exploit these vulnerabilities, thereby enhancing treatment efficacy and reducing the likelihood of resistance development. However, studies on collateral sensitivity usually involve laboratory-evolved isolates and the clinical relevance of the obtained mutations is not always clear. Further research is required to translate these findings into clinically applicable strategies.

An often-neglected aspect of laboratory evolution experiments is the impact of population dynamics on the experimental outcomes. Population composition is highly dependent on the population size within the experiment, the different frequencies of mutations that arise, and their phenotypic impact on resistance levels and growth fitness. Changes in experimental conditions can cause a specific phenotype to develop after 33,000 generations (Blount, et al. 2012; Good, et al. 2017) or after as few as 12 generations (Van Hofwegen, et al. 2016). The development of antibiotic resistance is another example in which experimental conditions profoundly impact the outcome (Huseby, et al. 2017; Garoff, et al. 2020). This is due to the multiple mechanisms by which chromosomal mutations can decrease antibiotic susceptibility. Resistance can be achieved through specific amino acid substitutions that alter drug-binding affinity or gene inactivation mutations that for example lead to increased drug efflux (Johanson and Hughes 1994; Sandegren and Andersson 2009; Fernandez and Hancock 2012). Gene inactivation mutations are approximately 1,000-fold more frequent (Schaaper, et al. 1986; Drake 1991; Abdulkarim and Hughes 1996). In smaller populations, these high-frequency mutations are overrepresented, and growth fitness is less important because of the lack of growth competition. As the population size increases, rare mutations appear and competitive fitness becomes important (Huseby, et al. 2017; Garoff, et al. 2020). Experimental studies to detect collateral sensitives are often performed in small volumes to enable high-throughput screening (Barbosa, et al. 2017; Imamovic, et al. 2018; Carolus, et al. 2024). Thus, these studies might suffer from a mutational bias that favours specific high-frequency mutations, which might not be clinically relevant.

In this study, we explored the clinical relevance of collateral sensitivity in six critical bacterial pathogens (Rice 2008). Utilizing a combination of experimental evolution, whole genome sequencing, genetic engineering, and phenotypic assays, we *(i)* engineered *E. coli* strains with clinically relevant resistance mutations to evaluate their susceptibility to various antibiotics, and *(ii)* investigated the evolutionary conditions under which collateral sensitivity arises in six bacterial species. Our findings revealed that in *E. coli*, collateral sensitivity was not associated with clinically relevant mutations but was instead linked to the reduced or abolished activity of the genes *guaA*, *metG*, *mnmA*, *sspA*, and *tusC*. Mutations in these genes generally incurred a significant fitness cost and were absent in ciprofloxacin-resistant clinical isolates of *E. coli*. Furthermore, we observed distinct collateral effects that emerged during ciprofloxacin resistance evolution in five out of six bacterial species, with *P. aeruginosa* showing the most conserved development of collateral sensitivity. These results underscore the challenges that must be addressed when developing collateral-sensitivity-based treatment strategies and highlight *P. aeruginosa* as a potential candidate for such treatments.

## Results and Discussion

### Clinically relevant resistance mutations do not result in collateral sensitivity in *E. coli*

Collateral sensitivity is often observed as a consequence of antibiotic resistance development during laboratory evolution experiments (Gonzales, et al. 2015; Barbosa, et al. 2017; Imamovic, et al. 2018; Podnecky, et al. 2018; Trigos, et al. 2021; Hasan, et al. 2023; Carolus, et al. 2024). These experiments yield complex genotypes and do not necessarily select for clinically relevant resistance mutations (Huseby, et al. 2017; Garoff, et al. 2020). We constructed a set of 13 *Escherichia coli* MG1655 strains harboring chromosomal resistance mutations frequently identified in clinical isolates of *E. coli, P. aeruginosa*, *Staphylococcus aureus*, and *M. tuberculosis* (Ma, et al. 1996; Sreevatsan, et al. 1996; Aubry-Damon, et al. 1998; Hooper 1999; Lee, et al. 2005; Sandegren, et al. 2008; Gygli, et al. 2019). These mutations include amino acid substitutions in the genes *gyrA* and *parC* (fluoroquinolone resistance), *rpsL* (streptomycin resistance), and *rpoB* (rifampicin resistance). Additionally, deletions of the genes *nfsA* and *nfsB* (nitrofurantoin resistance) as well as *acrR*, *marR*, and *soxR* (regulators of the multi-drug eflux pump AcrAB-TolC) were constructed. The antibiotic susceptibility of these 13 strains was assessed against 23 antibiotics from 17 distinct classes (Table 1). The results indicated that resistance and cross-resistance were common, with a total of 38 instances of decreased antibiotic susceptibility (Fig. 1A). In contrast, only two cases of weak collateral sensitivity were detected, both in the strain with the *rpoB*(Ser531Leu) mutation.

**Table 1.** Antibiotics included in this study.

| Inhibits/causes |  | Classification |  | Antibiotic | Abbreviation |
| --- | --- | --- | --- | --- | --- |
| Cell wall synthesis |  | Beta lactams | Penicillins | Ampicillin | AMP |
|  |  |  | Cephalosporins | Ceftaroline | CPT |
|  |  |  | Carbapenem | Meropenem | MEM |
|  |  |  | Monobactams | Aztreonam | ATM |
|  |  | Glycopeptides |  | Polymyxin B | PMB |
| Protein Synthesis | 30S | Aminoglycosides |  | Gentamicin | GEN |
|  |  |  |  | Streptomycin | STR |
|  |  |  |  | Amikacin | AMK |
|  |  | Aminocyclitols |  | Spectinomycin | SPT |
|  |  | Tetracyclines |  | Tetracyclin | TET |
|  |  |  |  | Tigecyclin | TGC |
|  | 50S | Oxazolidonones |  | Linezolid | LZD |
|  |  | Chloramphenicol |  | Chloramphenicol | CHL |
|  |  | Macrolides |  | Erythromycin | ERY |
|  |  | Lincosamides |  | Clindamycin | CLI |
|  |  | Diterpines |  | Tiamulin | TIA |
| DNA Topoisomerases |  | Quinolones |  | Nalidixic Acid | NAL |
|  |  | Fluoroquinolones |  | Ciprofloxacin | CIP |
| Folic Acid Synthesis |  | Sulfonamides |  | Sulfamethoxazole | SMX |
|  |  | DHFR Inhibitors |  | Trimethoprim | TMP |
| DNA damage |  | Metronidazole |  | Metronidazole | MTZ |
| RNA synthesis |  | Rifamicins |  | Rifampicin | RIF |
| Other |  | Nitrofurans |  | Nitrofurantoin | NIT |

**Fig. 1:**
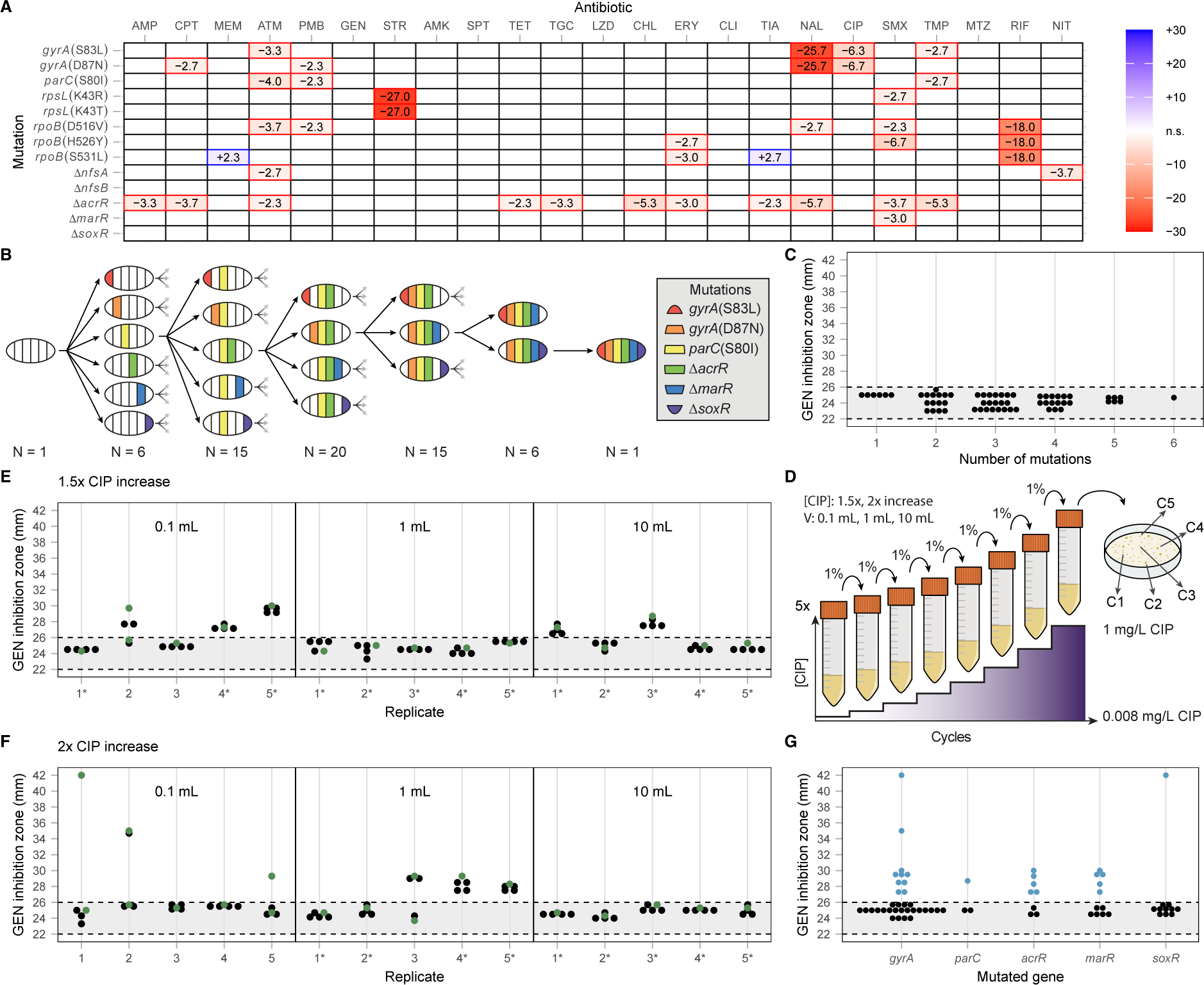
Collateral sensitivity to gentamicin is not caused by clinically relevant resistance mutations in *E. coli*. (**A**) Change in the zone of inhibition between wild-type *E. coli* and isolates carrying single resistance mutations. Increased antibiotic resistance is indicated in red, and collateral sensitivity is shown in blue. Empty fields represent no significant change. (**B**) Schematic overview of the construction of 64 strains with combinations of clinically relevant ciprofloxacin resistance mutations and (**C**) their respective gentamicin inhibition zones. (**D**) Schematic overview of the evolution experiment to select ciprofloxacin-resistant isolates. (**E**, **F**) Gentamicin inhibition zones of the 150 isolates from the evolution experiments with (**E**) 1.5-fold and (**F**) 2-fold increases in ciprofloxacin concentration per cycle. Evolution volumes are indicated within each graph, and lineages that survived until the final ciprofloxacin concentration of 1 mg/L are marked with an asterisk (*). Green dots represent isolates selected for whole genome sequencing. (**G**) Gentamicin zone of inhibition of whole genome sequenced isolates carrying mutations in the *gyrA*, *parC*, *acrR*, *marR*, and/or *soxR* genes. All values within the figure are averages of three biological replicates. The grey area between dotted lines represents the region not significantly different from the wild-type zone of inhibition. See Supplementary Data S1 for all individual measurements.

Previous studies have shown that resistance development to ciprofloxacin often results in collateral sensitivity to aminoglycoside antibiotics, which was not observed in our strains with single resistance mutations (Barbosa, et al. 2017; Podnecky, et al. 2018). In clinical *E. coli* isolates, high-level ciprofloxacin resistance is typically due to multiple mutations within the drug target genes (*gyrA* and *parC*) and efflux regulatory genes (*acrR*, *marR*, and *soxR*) (Hooper 1999; Huseby, et al. 2017). Thus, a combination of mutations might be required for the collateral sensitivity phenotype. To investigate this further, we constructed a set of 64 *E. coli* strains with every possible combination of the six clinically relevant resistance mutations *gyrA*(Ser83Leu), *gyrA*(Asp87Asn), *parC*(Ser80Ile), Δ*acrR*, Δ*marR*, and Δ*soxR* and measured their susceptibility to the aminoglycoside antibiotic gentamicin (Figs. 1B and C). None of the 64 constructed strains exhibited detectable collateral sensitivity, suggesting that the previously observed effects may not be attributable to the clinically relevant resistance mutations tested in this study.

### Collateral sensitivity develops during ciprofloxacin resistance evolution in *E. coli*

Collateral sensitivity development towards aminoglycoside antibiotics was not detected in any of the 64 strains with combinations of clinically relevant ciprofloxacin resistance mutations. To obtain mutations not included in the strain construction, we evolved *E. coli* isolates to become ciprofloxacin resistant. Previous studies have shown that differences in selective conditions significantly impact the outcome of ciprofloxacin resistance development (Garoff, et al. 2020). Therefore, we conducted our evolution experiment under six distinct conditions. Three different culture volumes (0.1 mL, 1 mL, and 10 mL) were used to achieve varying population sizes and ciprofloxacin concentrations were increased either 1.5-fold or 2-fold per cycle. Ciprofloxacin concentrations started at 0.008 mg/L, corresponding to 0.5× MIC^CIP^ of wild-type *E. coli* MG1655, and increased each cycle to a final concentration of 1 mg/L, the clinical breakpoint for ciprofloxacin resistance in *E. coli* (Fig. 1D) (Van, et al. 2019). Each of the six selective conditions was performed in five biological replicates, resulting in 30 lineages.

The clinical breakpoint was reached in 22 out of the 30 lineages and was more likely to be achieved with larger population sizes and lower drug selection pressure (Supplementary Fig. S1). At the endpoint of the evolution (1 mg/L or the highest concentration with visible growth), five colonies were isolated from each lineage, resulting in a total of 150 strains (Fig. 1D). The level of susceptibility to gentamicin was measured for these 150 isolates, and we found that 29% (43/150) displayed collateral sensitivity (Figs. 1E and F, Supplementary Table S1). Among the populations that reached the clinical breakpoint, the proportion of collateral sensitivity was highest for the selection in 0.1 mL with a 1.5-fold increase (66%, 2 out of 3 populations) and in the 1 mL volume with a 2-fold drug increase (50%, 2 out of 4 populations). In both cases, these volumes were the lowest where at least one population reached the final ciprofloxacin concentration of 1 mg/L. Overall, the five isolates from each evolutionary lineage displayed comparable gentamicin susceptibility levels in most populations (25 out of 30). In the remaining lineages (5 out of 30), the five isolates were divided into two groups: one group displayed collateral sensitivity, and the other did not (Figs. 1E and F, Supplementary Table S1). For lineages with one group, we selected a single isolate for further analysis, and for lineages with two groups, we selected one isolate from each group. In total, 35 strains were selected, of which 11 showed collateral sensitivity to gentamicin and 24 did not. The minimal inhibitory concentration for ciprofloxacin was determined for these selected isolates (Supplementary Table S1). No significant difference in the MIC^CIP^ value was detected between isolates that displayed collateral sensitivity and those that did not (P = 0.16, Mann-Whitney U test).

Taken together, these results show that collateral sensitivity towards gentamicin developed frequently during ciprofloxacin resistance evolution, and the final ciprofloxacin resistance level was indistinguishable between isolates that developed collateral sensitivity and those that did not.

### Collateral sensitivity to aminoglycosides in *E. coli* is caused by mutations in the genes *guaA*, *metG*, *mnmA*, *sspA*, and *tusC*

To elucidate the genetic basis of collateral sensitivity to gentamicin, we performed whole genome sequencing on the 35 selected isolates (Supplementary Table S2). The sequenced isolates had an average of 3.5 genetic changes, with strains displaying collateral sensitivity carrying significantly more genetic changes compared to those without collateral sensitivity (4.3 vs. 3.2 changes, P = 0.00625, Mann-Whitney U test). Each isolate carried 1 – 3 mutations within the clinically relevant genes *gyrA*, *parC*, *acrR*, *marR*, and *soxR*. However, no correlation was observed between mutations in these genes and the development of collateral sensitivity (Fig. 1G), confirming our initial findings that collateral sensitivity to gentamicin is not caused by mutations within these clinically relevant resistance genes. One strain (strain GB281) with collateral sensitivity carried a 414 kb duplication on the chromosome and was otherwise genetically identical to another isolate (strain GB99) from the same lineage that did not display collateral sensitivity. This strain was excluded from further analysis due to the complexity of dissecting which part of the duplication caused the observed collateral sensitivity. Excluding the five clinically relevant genes (*gyrA*, *parC*, *acrR*, *marR*, and *soxR*) from the analysis, we found that the remaining 10 isolates displaying collateral sensitivity harbored 1 – 2 additional mutations not found in isolates without detectable collateral sensitivity. These mutations were identified in the genes *bipA*, *cmoB*, *gadC*, *guaA*, *metG*, *mnmA*, *mtlD*, *rluF*, *sspA*, *tusC*, and *yheO* (Fig. 2A).

**Fig. 2:**
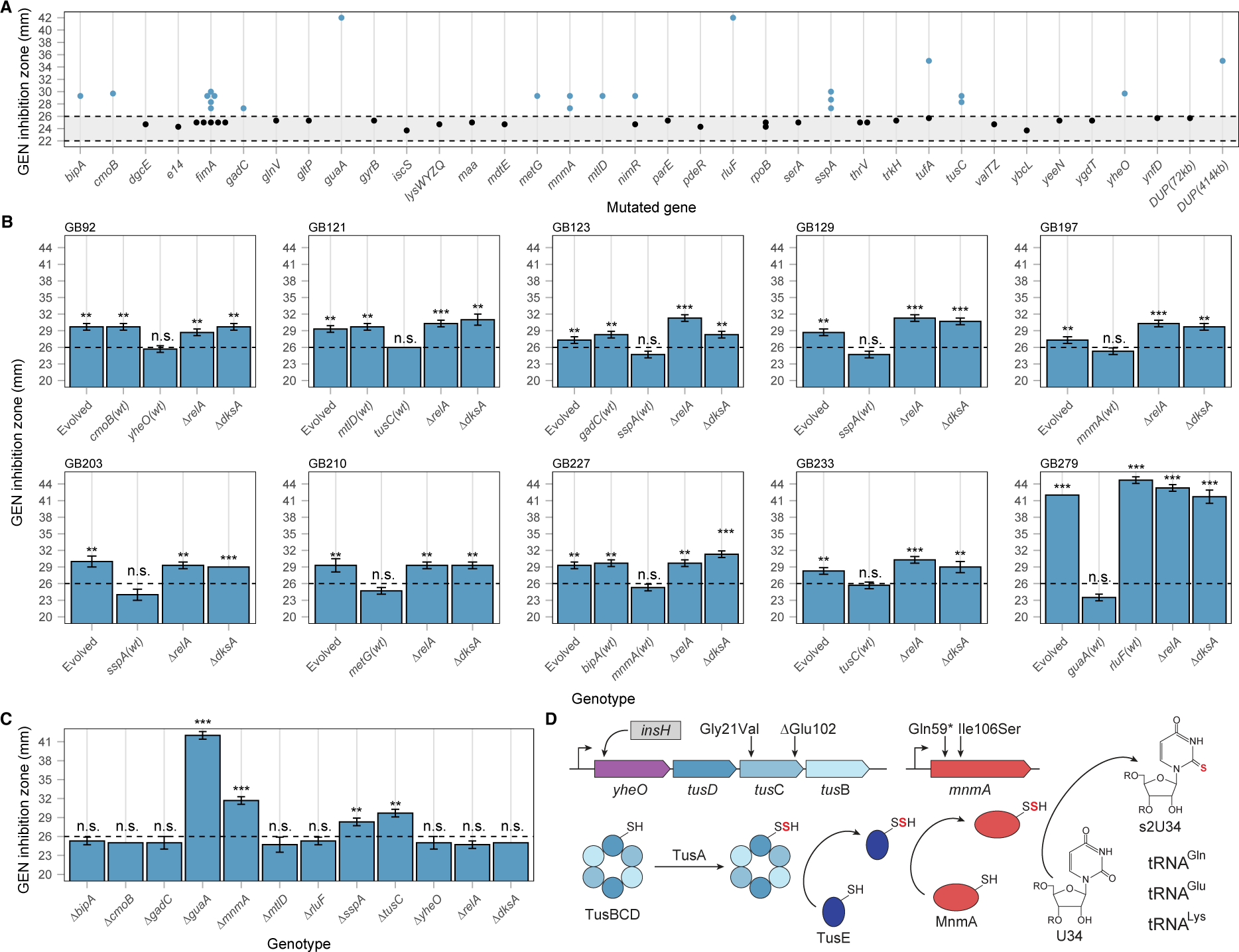
Collateral sensitivity to gentamicin in *E. coli* is caused by reduced function or inactivation of the genes *guaA*, *metG*, *mnmA*, *sspA*, and *tusC*. (**A**) Gentamicin zone of inhibition for whole genome sequenced isolates carrying mutations in the specified genes. The grey area between dotted lines represents the region not significantly different from the wild-type zone of inhibition. (**B**) Gentamicin zone of inhibition for evolved isolates displaying collateral sensitivity and the respective constructed strains with reintroduced wild-type alleles of specified genes or deletion of the genes *relA* and *dksA*. Each plot represents a specific evolved isolate, with the strain number stated above the plot. (**C**) Gentamicin zone of inhibition for strains with constructed single gene deletions. (**D**) Schematic overview of the *yheO*-*tusDCB* operon and the *mnmA* gene, showing the location of the identified mutations. TusBCD and MnmA are involved in the transfer of a sulfur (red) to the U34 residue of specific tRNAs. All values within the figure are averages of three biological replicates. See Supplementary Data S1 for all individual measurement values. The dotted line in the box plots indicates the minimal inhibition zone considered significantly different from the wild-type, and the error bars represent standard deviations. n.s.: not significant, **: P < 0.01, ***: P < 0.001.

To determine which mutations were responsible for the development of collateral sensitivity, we reintroduced the wild-type alleles of each of these 11 genes into the evolved isolates. No change in collateral sensitivity was observed when the mutant alleles of *bipA*, *cmoB*, *gadC*, *mtlD*, and *rluF* were replaced with their respective wild-type alleles. However, removal of the mutations in *guaA*, *metG*, *mnmA*, *sspA*, *tusC*, and *yheO* resulted in the loss of the previously observed collateral sensitivity to gentamicin (Fig. 2B). Each of the ten isolates with a collateral-sensitivity phenotype carried exactly one mutation within one of these six genes. In three of these genes (*mnmA*, *sspA*, and *yheO*), the identified mutations were expected to lead to a loss of protein function (nonsense mutations, frameshift mutations, or the insertion of an IS element). To elucidate whether the genes linked with collateral sensitivity to gentamicin do so due to loss of protein function, we constructed wild-type *E. coli* strains with deletions of *guaA*, *mnmA*, *sspA*, *tusC*, and *yheO* (*metG* is essential for growth and was excluded for this test). We found that deletions of *guaA*, *mnmA*, *sspA*, and *tusC* resulted in collateral sensitivity to gentamicin, while deletion of *yheO* did not (Fig. 2C). The *yheO* gene is the first gene within the *yheO*-*tusDCB* operon, indicating that the insertion of an IS element within the *yheO* gene causes collateral sensitivity to gentamicin due to polar effects on the expression of the *tusDCB* genes rather than inactivation of the YheO protein itself (Fig. 2D). We also constructed deletions of *bipA*, *cmoB*, *gadC*, *mtlD*, and *rluF* to further validate that inactivation of these genes does not cause collateral sensitivity. As expected, none of the deletions resulted in increased sensitivity to gentamicin (Fig. 2C).

Resistance to aminoglycosides in *E. coli* is typically due to the acquisition of horizontally transferred resistance genes (Ramirez, et al. 2013; Cazares, et al. 2020; Lund, et al. 2023). To investigate whether collateral sensitivity could re-sensitize gentamicin-resistant isolates, we introduced deletions of the *guaA*, *mnmA*, *sspA*, and *tusC* genes into an *E. coli* strain harboring the resistance gene *aaaC1* from the transposon Tn21. The aaaC1 gene encodes a 3-N-acyltransferase that inactivates gentamicin (Ramirez and Tolmasky 2010). Deletion of *mnmA* and *tusC* did not alter the level of gentamicin resistance. However, inactivation of either *guaA* or *sspA* increased gentamicin sensitivity in the constructed isolate (Fig. 3C). Notably, deletion of the *guaA* gene resulted in a strain with gentamicin sensitivity comparable to that of wild-type *E. coli*. Thus, the genome sequencing and mutation analysis indicate that inactivation or reduced function of the *guaA*, *metG*, *mnmA*, *sspA*, and *tusC* genes lead to collateral sensitivity to gentamicin in *E. coli*, with mutations in *sspA* and *guaA* effectively re-sensitizing a gentamicin-resistant *E. coli* isolate.

**Fig. 3:**
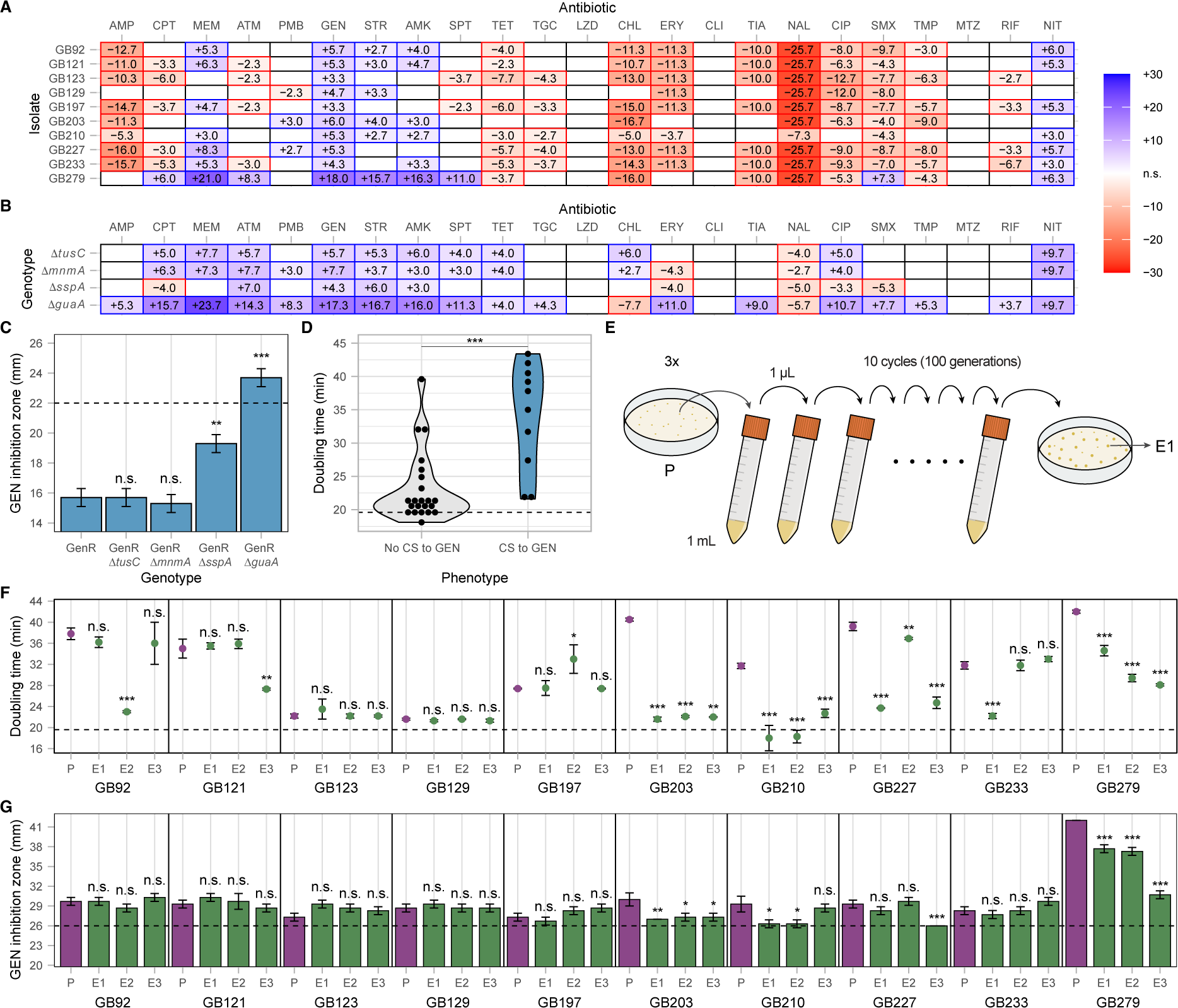
Ciprofloxacin-resistant isolates display complex antibiotic sensitivity and resistance profiles. **(A)** Change in the inhibition zone between wild-type *E. coli* and ciprofloxacin-resistant isolates displaying collateral sensitivity to gentamicin. (**B**) Change in the zone of inhibition between wild-type *E. coli* and constructed isolates with clean gene deletions. Increased antibiotic resistance is indicated in red, and collateral sensitivity is shown in blue. Empty fields represent no significant change. (**C**) Gentamicin zone of inhibition for *E. coli* strains carrying the gentamicin resistance gene *aaaC1* (GenR) and gene deletions causing collateral sensitivity to gentamicin. (**D**) Violin plot of exponential doubling times of ciprofloxacin-resistant isolates that do or do not display collateral sensitivity to gentamicin. (**E**) Schematic overview of the compensatory evolution experiment. (**F**) Exponential doubling times of parental strains and isolates after compensatory evolution. The dotted line indicates the doubling time of wild-type *E. coli*. (**G**) Gentamicin zone of inhibition for parental strains and isolates after compensatory evolution. The dotted line indicates the minimal inhibition zone considered significantly different from the wild-type *E. coli*. Evolved isolates (E1 - E3) are shown in green, and parental strains (P) in purple. All values within the figure are averages of three biological replicates, and the error bars represent standard deviations. See Supplementary Data S1 for all individual measurement values. n.s.: not significant, *: P < 0.05, **: P < 0.01, ***: P < 0.001.

### The stringent response is not involved in the development of collateral sensitivity to gentamicin in *E. coli*

One of the identified mutations causing collateral sensitivity is located within the aminoacyl-tRNA synthetase gene *metG*. The mutation *metG*(Phe305Cys) is situated in a protein region where mutations can lead to decreased methionine affinity and a diminished ability to discriminate against homocysteine (Fourmy, et al. 1991; Kim, et al. 1993). Additionally, mutations in the genes *tusC* and *mnmA* are expected to reduce or abolish the s^2^U^34^ modification in tRNA^Gln^, tRNA^Glu^, and tRNA^Lys^ (Fig. 2D), which is crucial for efficient tRNA charging (Giege, et al. 1998; Madore, et al. 1999). Previous studies have shown that mutations that reduce tRNA charging efficiency decrease susceptibility to ciprofloxacin through RelA-dependent activation of the stringent response (Wendrich, et al. 2002; Garoff, et al. 2018). To test if collateral sensitivity to gentamicin is also caused by activation of the stringent response, we deleted the *relA* gene in the ten evolved isolates displaying collateral sensitivity and in wild-type strains with clean gene deletions of *guaA*, *mnmA*, *sspA*, and *tusC*. Additionally, we deleted the transcription factor DksA, responsible for transcriptional reprogramming upon activation of the stringent response, within the ten evolved isolates (Doniselli, et al. 2015; Molodtsov, et al. 2018). Deletion of *relA* or *dksA* in wild-type *E. coli* did not change susceptibility to gentamicin, indicating that these individual deletions do not cause resistance or collateral sensitivity to gentamicin by themselves (Fig. 2C). None of the constructed strains displayed diminished collateral sensitivity to gentamicin when the genes were deleted in the evolved isolates (Fig. 2B). Similarly, combining deletions of *guaA*, *mnmA*, *sspA*, or *tusC* with deletions of *relA* did not result in loss of collateral sensitivity (Supplementary Fig. S2). These results indicate that the stringent response is not involved in the development of collateral sensitivity to gentamicin.

### Mutations in *guaA*, *metG*, *mnmA*, *sspA*, and *tusC* cause complex collateral sensitivity and resistance profiles

To extend our analysis, we measured the change in drug susceptibility of the 10 ciprofloxacin-resistant isolates that display collateral sensitivity to gentamicin across the set of 23 antibiotics included in this study (Table 1). Increased resistance to eleven antibiotics (AMP, CPT, TET, TGC, CHL, ERY, TIA, NAL, CIP, SMX, and TMP) and collateral sensitivity to five antibiotics (MEM, GEN, STR, AMK, and NIT) were observed in at least half of the isolates. In total, we identified 96 instances of increased resistance and 42 cases of collateral sensitivity (Fig. 3A). We also measured sensitivity levels to the 23 antibiotics in the evolved strains where mutations in the genes *bipA*, *cmoB*, *gadC*, *guaA*, *metG*, *mnmA*, *mtlD*, *rluF*, *sspA*, *tusC*, and *yheO* were replaced with their respective wild-type alleles (Supplementary Fig. S3). Consistent with previous results, only mutations in *guaA*, *metG*, *mnmA*, *sspA*, *tusC*, and *yheO* contributed to collateral sensitivity towards any antibiotic, while mutations in *bipA*, *cmoB*, *gadC*, *mtlD*, and *rluF* did not.

The complex genotypes of the evolved lineages complicate the dissection of specific mutation effects. Therefore, we measured antibiotic sensitivity in clean strains with deletions of *guaA*, *mnmA*, *sspA*, and *tusC* (Fig. 3B). The results show that the mutations fall into three distinct groups: *(i)* Δ*guaA* increases sensitivity to 18 antibiotics and resistance to 2 antibiotics, *(ii)* Δ*mnmA* and Δ*tusC* increase sensitivity to 11 antibiotics and resistance to 1 antibiotic, and (iii) Δ*sspA* increases sensitivity to 4 and resistance to 5 antibiotics. The distinct antibiotic sensitivity/resistance profiles of these groups suggest different underlying molecular mechanisms.

To test if the stringent response is involved in any of the observed sensitivity or resistance effects, we measured antibiotic sensitivity in the strains with additional deletion of the *relA* gene (Supplementary Fig. S2). Deleting *relA* generally increased antibiotic sensitivity and abolished 7 out of the 10 previously observed cases of increased antibiotic resistance. These data indicate that deletions of the *guaA*, *mnmA*, *sspA*, and *tusC* genes activate the stringent response, which decreases some of the collateral sensitivity affects caused by the gene deletions. However, the stringent response does not significantly contribute to any of the observed collateral sensitivity effects.

### No evidence for clinical relevance of mutations that cause collateral sensitivity in *E. coli*

Previous studies have shown that clinical ciprofloxacin resistance develops through a combination of mutations that do not reduce cellular growth fitness (Huseby, et al. 2017; Praski Alzrigat, et al. 2017). To investigate this, we measured the exponential growth rate of 35 representative isolates from this study (Fig. 3D). The results show that isolates displaying collateral sensitivity to gentamicin have a significantly longer exponential doubling time compared to those that do not exhibit collateral sensitivity (33.9 ± 7.6 min vs. 22.7 ± 5.3 min, P = 0.0002, Mann-Whitney U test). This suggests that the development of collateral sensitivity is accompanied by a significant fitness cost, which is likely to be counter-selected in clinical setting. Fitness costs caused by antibiotic resistance mutations can be rapidly ameliorated during compensatory evolution. We tested this by cycling the ten isolates that displayed collateral sensitivity in the absence of drug selection (Fig. 3E). After 100 generations of compensatory evolution, half of the evolved lineages (15/30) showed a significantly faster exponential doubling time compared to their respective unevolved parental strain (Fig. 3F). In 9 out of 15 cases (60%), the increase in exponential growth rate was accompanied by a reduction in or loss of collateral sensitivity to gentamicin, indicating that the collateral sensitivity phenotype is frequently lost during growth-compensatory adaptation (Fig. 3G). These data suggest that collateral sensitivity to gentamicin is associated with a significant fitness cost and is quickly lost when selective conditions favor high-fitness isolates, reducing the clinical relevance of these mutations. To determine if mutations within the *guaA*, *metG*, *mnmA*, *sspA*, and *tusC* genes are relevant during the evolution of ciprofloxacin resistance in clinical settings, we analyzed a set of 835 genomes from clinical *E. coli* isolates. Each genome was classified as likely ciprofloxacin-resistant or - sensitive based on the presence of mutations in the *gyrA* (amino acids Ser83 or Asp87) and *parC* (amino acids Ser80 and Glu84) genes. In total, 403 genomes were classified as ciprofloxacin-sensitive and 432 as ciprofloxacin-resistant (Table S3). A comparison of the *guaA*, *metG*, *mnmA*, *sspA*, and *tusC* genes within these two groups showed no significant increase in mutations within the ciprofloxacin-resistant group (Table S4). Thus, we find no evidence that mutations that cause collateral sensitivity are selected for in clinical *E. coli* isolates during ciprofloxacin resistance evolution.

### Critical bacterial pathogens exhibit distinct collateral responses during ciprofloxacin resistance evolution

The analysis of *E. coli* suggests that evolutionary trajectories leading to collateral sensitivity are unlikely to be clinically relevant. To determine if this holds true for other species, we evolved ciprofloxacin resistance in five additional critical bacterial pathogens: *Salmonella Typhimurium*, *Klebsiella pneumoniae*, *Acinetobacter baumanii*, *P. aeruginosa*, and *S. aureus* (Fig. 4F). We measured gentamicin resistance levels in endpoint isolates and observed distinct collateral responses among the species (Figs. 4A-E). For *K. pneumoniae*, none of the isolates exhibited changes in gentamicin sensitivity compared to the parental wild-type strain. A small minority of *S. Typhimurium* (2 lineages) and *S. aureus* (1 lineage) isolates displayed collateral sensitivity, which was only observed in the smallest evolution volume. Notably, collateral sensitivity was most prevalent in the evolved *P. aeruginosa* strains, with isolates in 8 out of 9 lineages showing increased sensitivity to gentamicin. In the evolved *A. baumanii* strains, collateral sensitivity to gentamicin was absent, although 6 out of 9 lineages exhibited collateral resistance to gentamicin.

**Fig. 4:**
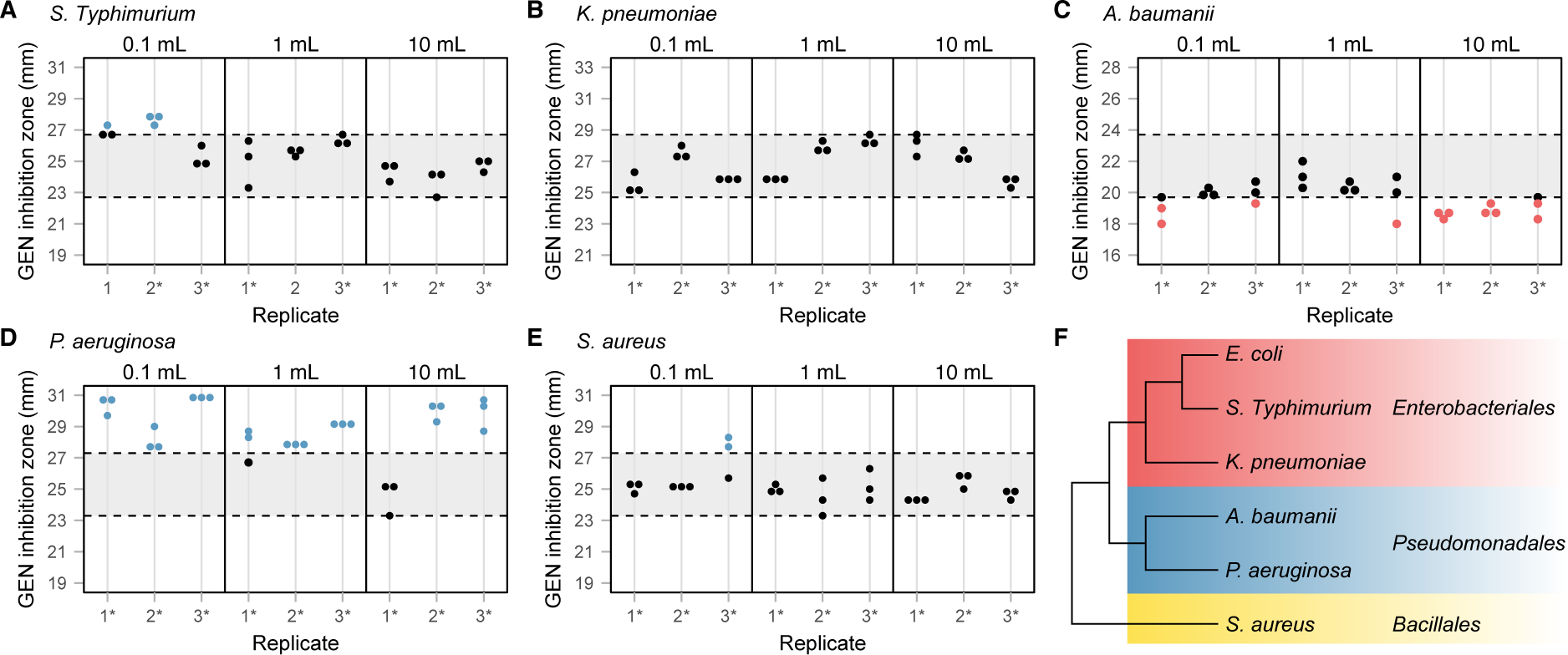
Critical bacterial pathogens exhibit distinct collateral responses during ciprofloxacin resistance evolution. (**A**-**E**) Gentamicin inhibition zones for ciprofloxacin-resistant isolates of (**A**) *S. Typhimurium*, (**B**) *K. pneumoniae*, (**C**) *A. baumanii*, (**D**) *P. aeruginosa*, and (**E**) *S. aureus*. Evolution volumes are indicated above each graph, and replicates that survived until the final ciprofloxacin concentration of 1 mg/L are marked with an asterisk (*). Blue dots represent isolates displaying collateral sensitivity and red dots represent isolates displaying collateral resistance. All values are averages of three biological replicates. The grey area between dotted lines represents the region not significantly different from the respective wild-type zone of inhibition. See Supplementary Data S1 for all individual measurement values. (**F**) Phylogenetic relationship of the species included in this study(Brandis 2021).

These data demonstrate that different bacterial species exhibit distinct collateral responses during antibiotic resistance development. All six species in this study displayed multiple evolutionary trajectories during ciprofloxacin resistance development, with some trajectories not resulting in collateral effects to gentamicin, while others led to increased gentamicin sensitivity or resistance. The consistent development of collateral sensitivity in *P. aeruginosa* isolates across all tested evolution volumes suggests that major evolutionary trajectories in this species lead to collateral effects. This highlights *P. aeruginosa* as a promising candidate for collateral-sensitivity-based treatment strategies.

## Conclusions

Our study demonstrates that collateral sensitivity to gentamicin in *E. coli* arises from mutations in the *tusC*, *mnmA*, *metG*, *sspA*, or *guaA* genes, which impair or eliminate their function. Previous research has indicated that mutations in these or related genes are selected during the development of ciprofloxacin resistance. These mutations either activate the stringent response or mimic binding ppGpp binding to the RNA polymerase, leading to transcriptional changes that enhance ciprofloxacin efflux (Garoff, et al. 2018; Brandis, et al. 2021). However, our findings reveal that collateral sensitivity to gentamicin is independent of the stringent response.

The antibiotic resistance profiles of isolates with clean gene deletions suggest that the mechanisms underlying gentamicin sensitivity fall into at least three distinct groups (excluding *metG*, which cannot be deleted). The most common group includes mutations affecting *tusC* and *mnmA*, found in 5 out of 10 evolved isolates. The modification of U^34^ by MnmA is essential for efficient tRNA charging (Giege, et al. 1998; Madore, et al. 1999). Thus, deletion of the *tusC* or *mnmA* likely activates the stringent response, contributing to ciprofloxacin resistance. Additionally, deletion of *mnmA* increases the translational error rate due to undermodified tRNAs (Urbonavicius, et al. 2001; Manickam, et al. 2016), making isolates more sensitive to gentamicin as lower antibiotic concentrations are required to reach lethal translational error rates (Fig 5).

**Fig. 5:**
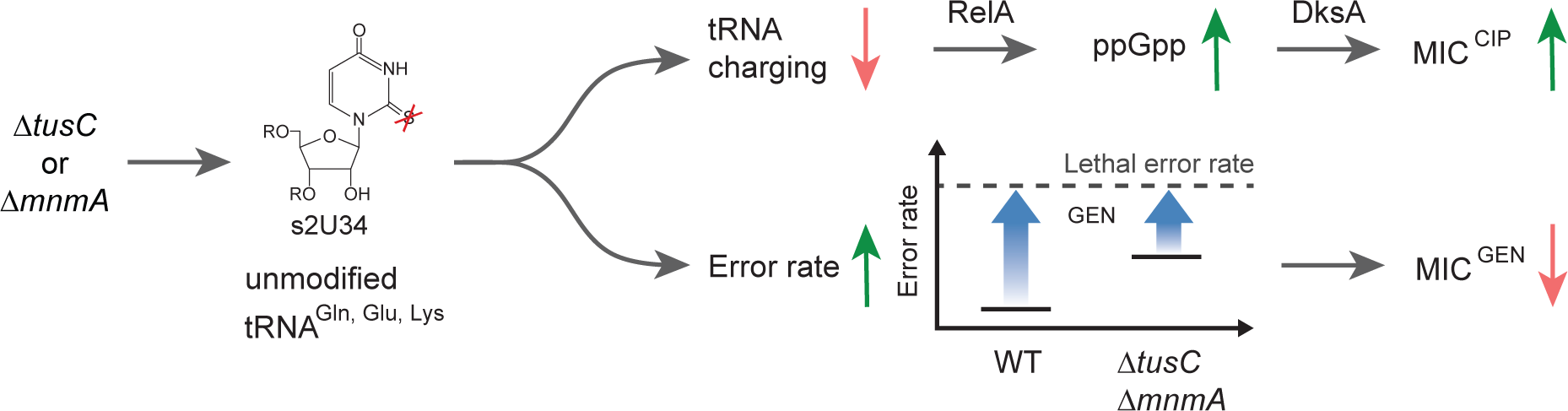
Proposed molecular mechanism underlying collateral sensitivity to gentamicin in *E. coli* with *tusC* or *mnmA* gene deletion. Deletion of the *tusC* or *mnmA* gene abolished s^2^U^34^ modification of specific tRNAs. This undermodification results in less efficient charging of tRNAs, leading to activation of the stringent response via RelA/DksA. Consequently, this activation increases drug eflux, resulting in heightened resistance to ciprofloxacin (top). Additionally, the absence of s^2^U^34^ tRNA modifications elevates the basal translational error rate in the isolates, causing collateral sensitivity to gentamicin (bottom).

Our experiments show that collateral sensitivity in *E. coli* can develop during resistance evolution and that it can re-sensitize isolates with horizontally transferred resistance genes. However, several caveats regarding the clinical applicability of collateral-sensitivity-based treatment strategies for *E. coli* infections were identified: *(i)* Collateral sensitivity did not develop uniformly across all lineages and was absent in some selective conditions. *(ii)* The mutations causing collateral sensitivity are not associated with clinical resistance, and we did not detect these mutations in clinical *E. coli* isolates. *(iii)* Collateral sensitivity to gentamicin incurred a significant fitness cost and was rapidly lost during compensatory evolution.

These results underscore the challenges of implementing collateral-sensitivity-based treatment strategies in clinical settings and highlight the importance of testing multiple evolutionary conditions and fitness effects when investigating novel phenotypes. Multiple evolutionary trajectories can lead to the selected phenotype, but only a subset will provide the desired secondary effects. For collateral sensitivity to be clinically relevant, the collateral effect must result from a mutation on a clinically relevant trajectory. Among the six species tested in this study, *P. aeruginosa* emerged as the most promising candidate for such a treatment strategy.

## Methods

### Bacterial strains and growth conditions

All strains were derived from *Escherichia coli* K-12 strain MG1655 (Blattner, et al. 1997), *Salmonella enterica* serovar Typhimurium strain 14028s (Jarvik, et al. 2010), *Pseudomonas aeruginosa* strain PAO1 (Stover, et al. 2000), *Klebsiella pneumoniae* strain ATCC 13883, *Acinetobacter baumanii* strain ATCC 19606, or *Staphylococcus aureus* strain ATCC 29213. Bacteria were generally grown at 37 °C in Luria Broth (LB; 10 g/L tryptone, 5 g/L yeast extract, 5 g/L NaCl) or on LB agar plates (LA; LB solidified with 1.5% Oxoid agar). Strains containing temperature-sensitive pSIM plasmids were maintained at 30 °C (Datta, et al. 2006; Koskiniemi, et al. 2011). Chloramphenicol (25 mg/L or 50 mg/L), rifampicin (100 mg/L), streptomycin (100 mg/L), tetracycline (15 mg/L), and sucrose (50 g/L) were added to the media as required. Ciprofloxacin was added at various concentrations during the evolution experiments.

### Genetic engineering

Mutations in *gyrA*, *rpoB*, and *rpsL* were introduced into wild-type *E. coli* by lambda-red recombineering using ssDNA oligonucleotides (Ellis, et al. 2001), selecting for increased resistance to ciprofloxacin, rifampicin, and streptomycin, respectively. The *parC*(Ser80Ile) mutation was introduced into *E. coli* with *gyrA*(Ser83Leu + Asp87Asn) mutations by lambda-red recombineering using ssDNA oligonucleotides, selecting for increased ciprofloxacin resistance. Gene deletions were constructed using DiRex(Nasvall 2017). All deletions were designed to omit the first six and last six amino acids, and no known regulatory elements for unrelated genes were deleted. Mutant alleles were moved between strains by P1 transduction using Dup-In or direct selection for DiRex intermediates of deletions (Ikeda and Tomizawa 1965; Nasvall, et al. 2017).

### PCR and DNA sequencing

DNA amplifications were performed using 2× Taq PCR Mastermix (Thermo Scientific) or 2× Phusion PCR Mastermix (New England Biolabs) according to the manufacturer’s protocol. Genomic DNA was prepared using the DNeasy UltraClean Microbial Kit (Qiagen) following the manufacturer’s instructions. Sanger sequencing of PCR products and whole genome sequencing was performed by Macrogen (The Netherlands). Sequences were analyzed using the CLC Genomics Workbench 24.0.1 (CLCbio, Qiagen).

### Antibiotic susceptibility testing

Minimal inhibitory concentration (MIC) testing was conducted following EUCAST guidelines by broth microdilution in LB media, with incubation for 18 – 20 h at 37 °C. Collateral sensitivity was determined using the disc diffusion method. Colonies were resuspended in 0.9% NaCl solution to 0.5 MacFarland standard. Bacteria were spread on LA plates using sterile cotton swaps, antibiotic disc were applied, and plates were incubated for 18 – 20 h at 37 °C. A list of all tested antibiotic and the quantity of antibiotic in each disc is provided in Supplementary Table S5. Tests were performed in biological triplicates, and changes in the zone of inhibition were considered significant if the mean change was >2 mm and a two-sided t-test indicated significant differences.

### Evolution of resistance to ciprofloxacin

Evolution of resistance to ciprofloxacin was performed with three different volumes: 0.1 mL, 1 mL, and 10 mL. Cultures were grown in 15 mL Falcon tubes (0.1 mL and 1 mL) or 50 mL Falcon tubes (10 mL). Independent lineages were grown shaking at 37 °C in LB to initiate cultures. Every 24 h, cultures with visible growth were diluted 100-fold in fresh media. The concentration of ciprofloxacin in the media was stepwise increase from 0.008 mg/L (0.5× MIC of wild-type *E. coli* MG1655) to the clinical breakpoint of 1 mg/L using 1.5-fold or 2-fold increments (Supplementary Table S6). The culture at the highest ciprofloxacin concentration with observable growth (end-point culture) of each lineage was stocked at -80 °C in 15% glycerol. Each evolution was performed in five independent replicates (three replicates for species other than *E. coli*). End-point populations were diluted from the freezer into 0.9% NaCl solutions and plated onto LA plates to obtain single colonies. Five colonies were isolated from each end-point lineage for further analysis (three colonies for species other than *E. coli*).

### Genome analysis of clinical isolates

*E. coli* genomes were downloaded on the 27^th^ Aug 2024 from the Bacterial and Viral Bioinformatics Resource Center (BV-BRC) (Olson, et al. 2023) with the following filters. Genome Status: Complete, Genome Quality: Good, Host Group: Human. Genomes with a size below 4 Mb or a Sequencing Depth <100× were removed from the collection. To further ensure good sequencing quality, all genomes were analyzed for a set of essential genes (*dnaA*, *fusA*, *gyrA*, *parC*, *rplA*, *rpoA*, *rpoB*, *rpoC*, *rpsA*) with a combined length of >20 kb. Genomes containing frameshift mutations within any of these genes were excluded, resulting in a set of 835 genomes. All genomes were classified as ciprofloxacin-sensitive (CIP^S^) or ciprofloxacin-resistant (CIP^R^) based on the presence of mutations in *gyrA* (amino acids Ser83 and Asp87) and *parC* (amino acids Ser80 and Glu84). See Supplementary Table S3 for an overview of identified mutations. Protein sequences for all genes of interest were extracted from the genomes and aligned to determine the conservation of each amino acid within the proteins. Positions with conservation below 99% were ignored from the further analysis, and genomes with mutations in the remaining parts of the protein sequences were identified. All sequence analyses were performed using the CLC Genomics Workbench 24.0.1 (CLCbio, Qiagen). For each gene, the likelihood of mutations being present in ciprofloxacin-resistant isolates compared to sensitive ones was determined using a two-sided Fisher’s exact test (Supplementary Table S4).

### Growth rate measurements

Exponential growth rates were determined using a Bioscreen C Pro machine. Overnight cultures were 1,000-fold diluted in fresh LB and 300 µL of each culture were transferred into a Bioscreen Honeycomb 2 plate. Cultures were incubated for 18 h under continuous shaking and OD_600nm_ was measured every 5 min. Exponential growth rates were determined over a window of 10 measurement points (approximately 2 doubling of wild-type *E. coli*) from the point that the culture reaches an OD_600nm_ of 0.015. See Supplementary Data 2 for raw data.

## Acknowledgements

This work was supported by grants to GB from the Åke Wibergs foundation (grant numbers M22-0037 and M24-0011), the Tore Nilsons foundation (grant numbers 2022-004, 2023-083, and 2024-148), the Swedish Research Council (grant number 2023-03718), and the Carl Tryggers foundation (grant number CTS 22:1886).

## Conflict of Interest

The authors declare no conflict of interest.

## Data availability

The raw whole-genome sequencing data are available at the NCBI SRA database (SUB15311114). All raw data are provided with this paper in Supplementary Data S1 and S2. Strains used in this study are available upon request to the corresponding author.

## Author contributions

V.C., L.E., Y.C., C.D., E.C.D., T.I., G.M., E.P., A.S., E.Z., A.af.K., and A.K. performed experiments. V.C. and G.B. analyzed experimental data sets. G.B. conceived and designed the study and wrote the initial manuscript. All authors reviewed and edited the manuscript.

## References

Abdulkarim F, Hughes D. 1996. Homologous recombination between the tuf genes of Salmonella typhimurium. J Mol Biol 260:506–522.

Aubry-Damon H, Soussy CJ, Courvalin P. 1998. Characterization of mutations in the rpoB gene that confer rifampin resistance in Staphylococcus aureus. Antimicrob Agents Chemother 42:2590–2594.

Barbosa C, Trebosc V, Kemmer C, Rosenstiel P, Beardmore R, Schulenburg H, Jansen G. 2017. Alternative Evolutionary Paths to Bacterial Antibiotic Resistance Cause Distinct Collateral Effects. Mol Biol Evol 34:2229–2244.

Blattner FR, Plunkett G, 3rd, Bloch CA, Perna NT, Burland V, Riley M, Collado-Vides J, Glasner JD, Rode CK, Mayhew GF, et al. 1997. The complete genome sequence of Escherichia coli K-12. Science 277:1453–1462.

Blount ZD, Barrick JE, Davidson CJ, Lenski RE. 2012. Genomic analysis of a key innovation in an experimental Escherichia coli population. Nature 489:513–518.

Brandis G. 2021. Reconstructing the Evolutionary History of a Highly Conserved Operon Cluster in Gammaproteobacteria and Bacilli. Genome Biol Evol 13.

Brandis G, Granstrom S, Leber AT, Bartke K, Garoff L, Cao S, Huseby DL, Hughes D. 2021. Mutant RNA polymerase can reduce susceptibility to antibiotics via ppGpp-independent induction of a stringent-like response. J Antimicrob Chemother 76:606–615.

Carolus H, Sofras D, Boccarella G, Jacobs S, Biriukov V, Goossens L, Chen A, Vantyghem I, Verbeeck T, Pierson S, et al. 2024. Collateral sensitivity counteracts the evolution of antifungal drug resistance in Candida auris. Nat Microbiol 9:2954–2969.

Cazares A, Moore MP, Hall JPJ, Wright LL, Grimes M, Emond-Rheault JG, Pongchaikul P, Santanirand P, Levesque RC, Fothergill JL, et al. 2020. A megaplasmid family driving dissemination of multidrug resistance in Pseudomonas. Nat Commun 11:1370.

Datta S, Costantino N, Court DL. 2006. A set of recombineering plasmids for gram-negative bacteria. Gene 379:109–115.

Doniselli N, Rodriguez-Aliaga P, Amidani D, Bardales JA, Bustamante C, Guerra DG, Rivetti C. 2015. New insights into the regulatory mechanisms of ppGpp and DksA on Escherichia coli RNA polymerase-promoter complex. Nucleic Acids Res 43:5249–5262.

Drake JW. 1991. A constant rate of spontaneous mutation in DNA-based microbes. Proc Natl Acad Sci U S A 88:7160–7164.

Ellis HM, Yu D, DiTizio T, Court DL. 2001. High efficiency mutagenesis, repair, and engineering of chromosomal DNA using single-stranded oligonucleotides. Proc Natl Acad Sci U S A 98:6742–6746.

Fernandez L, Hancock RE. 2012. Adaptive and mutational resistance: role of porins and eflux pumps in drug resistance. Clin Microbiol Rev 25:661–681.

Fourmy D, Mechulam Y, Brunie S, Blanquet S, Fayat G. 1991. Identification of residues involved in the binding of methionine by Escherichia coli methionyl-tRNA synthetase. FEBS Lett 292:259–263.

Garoff L, Huseby DL, Praski Alzrigat L, Hughes D. 2018. Effect of aminoacyl-tRNA synthetase mutations on susceptibility to ciprofloxacin in Escherichia coli. J Antimicrob Chemother 73:3285–3292.

Garoff L, Pietsch F, Huseby DL, Lilja T, Brandis G, Hughes D. 2020. Population Bottlenecks Strongly Influence the Evolutionary Trajectory to Fluoroquinolone Resistance in Escherichia coli. Mol Biol Evol 37:1637–1646.

GBD 2021 Antimicrobial Resistance Collaborators. 2024. Global burden of bacterial antimicrobial resistance 1990-2021: a systematic analysis with forecasts to 2050. Lancet 404:1199–1226.

Giege R, Sissler M, Florentz C. 1998. Universal rules and idiosyncratic features in tRNA identity. Nucleic Acids Res 26:5017–5035.

Gonzales PR, Pesesky MW, Bouley R, Ballard A, Biddy BA, Suckow MA, Wolter WR, Schroeder VA, Burnham CA, Mobashery S, et al. 2015. Synergistic, collaterally sensitive beta-lactam combinations suppress resistance in MRSA. Nat Chem Biol 11:855–861.

Good BH, McDonald MJ, Barrick JE, Lenski RE, Desai MM. 2017. The dynamics of molecular evolution over 60,000 generations. Nature 551:45–50.

Gygli SM, Keller PM, Ballif M, Blochliger N, Homke R, Reinhard M, Loiseau C, Ritter C, Sander P, Borrell S, et al. 2019. Whole-Genome Sequencing for Drug Resistance Profile Prediction in Mycobacterium tuberculosis. Antimicrob Agents Chemother 63.

Hasan M, Wang J, Ahn J. 2023. Ciprofloxacin and Tetracycline Resistance Cause Collateral Sensitivity to Aminoglycosides in Salmonella Typhimurium. Antibiotics (Basel) 12.

Hooper DC. 1999. Mechanisms of fluoroquinolone resistance. Drug Resist Updat 2:38–55.

Huseby DL, Pietsch F, Brandis G, Garoff L, Tegehall A, Hughes D. 2017. Mutation Supply and Relative Fitness Shape the Genotypes of Ciprofloxacin-Resistant Escherichia coli. Mol Biol Evol 34:1029–1039.

Ikeda H, Tomizawa JI. 1965. Transducing fragments in generalized transduction by phage P1. I. Molecular origin of the fragments. J Mol Biol 14:85–109.

Imamovic L, Ellabaan MMH, Dantas Machado AM, Citterio L, Wulff T, Molin S, Krogh Johansen H, Sommer MOA. 2018. Drug-Driven Phenotypic Convergence Supports Rational Treatment Strategies of Chronic Infections. Cell 172:121–134 e114.

Jarvik T, Smillie C, Groisman EA, Ochman H. 2010. Short-term signatures of evolutionary change in the Salmonella enterica serovar typhimurium 14028 genome. J Bacteriol 192:560–567.

Johanson U, Hughes D. 1994. Fusidic acid-resistant mutants define three regions in elongation factor G of Salmonella typhimurium. Gene 143:55–59.

Kim HY, Ghosh G, Schulman LH, Brunie S, Jakubowski H. 1993. The relationship between synthetic and editing functions of the active site of an aminoacyl-tRNA synthetase. Proc Natl Acad Sci U S A 90:11553–11557.

Koskiniemi S, Pranting M, Gullberg E, Nasvall J, Andersson DI. 2011. Activation of cryptic aminoglycoside resistance in Salmonella enterica. Mol Microbiol 80:1464–1478.

Lee JK, Lee YS, Park YK, Kim BS. 2005. Alterations in the GyrA and GyrB subunits of topoisomerase II and the ParC and ParE subunits of topoisomerase IV in ciprofloxacin-resistant clinical isolates of Pseudomonas aeruginosa. Int J Antimicrob Agents 25:290–295.

Lund D, Coertze RD, Parras-Molto M, Berglund F, Flach CF, Johnning A, Larsson DGJ, Kristiansson E. 2023. Extensive screening reveals previously undiscovered aminoglycoside resistance genes in human pathogens. Commun Biol 6:812.

Ma D, Alberti M, Lynch C, Nikaido H, Hearst JE. 1996. The local repressor AcrR plays a modulating role in the regulation of acrAB genes of Escherichia coli by global stress signals. Mol Microbiol 19:101–112.

Madore E, Florentz C, Giege R, Sekine S, Yokoyama S, Lapointe J. 1999. Effect of modified nucleotides on Escherichia coli tRNAGlu structure and on its aminoacylation by glutamyl-tRNA synthetase. Predominant and distinct roles of the mnm5 and s2 modifications of U34. Eur J Biochem 266:1128–1135.

Mahmud HA, Wakeman CA. 2024. Navigating collateral sensitivity: insights into the mechanisms and applications of antibiotic resistance trade-offs. Front Microbiol 15:1478789.

Maltas J, Huynh A, Wood KB. 2025. Dynamic collateral sensitivity profiles highlight opportunities and challenges for optimizing antibiotic treatments. PLoS Biol 23:e3002970.

Manickam N, Joshi K, Bhatt MJ, Farabaugh PJ. 2016. Effects of tRNA modification on translational accuracy depend on intrinsic codon-anticodon strength. Nucleic Acids Res 44:1871–1881.

Molodtsov V, Sineva E, Zhang L, Huang X, Cashel M, Ades SE, Murakami KS. 2018. Allosteric Effector ppGpp Potentiates the Inhibition of Transcript Initiation by DksA. Mol Cell 69:828–839 e825.

Mulet X, Moya B, Juan C, Macia MD, Perez JL, Blazquez J, Oliver A. 2011. Antagonistic interactions of Pseudomonas aeruginosa antibiotic resistance mechanisms in planktonic but not biofilm growth. Antimicrob Agents Chemother 55:4560–4568.

Nasvall J. 2017. Direct and Inverted Repeat stimulated excision (DIRex): Simple, single-step, and scar-free mutagenesis of bacterial genes. PLoS One 12:e0184126.

Nasvall J, Knoppel A, Andersson DI. 2017. Duplication-Insertion Recombineering: a fast and scar-free method for efficient transfer of multiple mutations in bacteria. Nucleic Acids Res 45:e33.

Olson RD, Assaf R, Brettin T, Conrad N, Cucinell C, Davis JJ, Dempsey DM, Dickerman A, Dietrich EM, Kenyon RW, et al. 2023. Introducing the Bacterial and Viral Bioinformatics Resource Center (BV-BRC): a resource combining PATRIC, IRD and ViPR. Nucleic Acids Res 51:D678–D689.

Podnecky NL, Fredheim EGA, Kloos J, Sorum V, Primicerio R, Roberts AP, Rozen DE, Samuelsen O, Johnsen PJ. 2018. Conserved collateral antibiotic susceptibility networks in diverse clinical strains of Escherichia coli. Nat Commun 9:3673.

Praski Alzrigat L, Huseby DL, Brandis G, Hughes D. 2017. Fitness cost constrains the spectrum of marR mutations in ciprofloxacin-resistant Escherichia coli. J Antimicrob Chemother 72:3016–3024.

Ramirez MS, Nikolaidis N, Tolmasky ME. 2013. Rise and dissemination of aminoglycoside resistance: the aac(6’)-Ib paradigm. Front Microbiol 4:121.

Ramirez MS, Tolmasky ME. 2010. Aminoglycoside modifying enzymes. Drug Resist Updat 13:151–171.

Rice LB. 2008. Federal funding for the study of antimicrobial resistance in nosocomial pathogens: no ESKAPE. J Infect Dis 197:1079–1081.

Roemhild R, Andersson DI. 2021. Mechanisms and therapeutic potential of collateral sensitivity to antibiotics. PLoS Pathog 17:e1009172.

Roemhild R, Linkevicius M, Andersson DI. 2020. Molecular mechanisms of collateral sensitivity to the antibiotic nitrofurantoin. PLoS Biol 18:e3000612.

Sandegren L, Andersson DI. 2009. Bacterial gene amplification: implications for the evolution of antibiotic resistance. Nat Rev Microbiol 7:578–588.

Sandegren L, Lindqvist A, Kahlmeter G, Andersson DI. 2008. Nitrofurantoin resistance mechanism and fitness cost in Escherichia coli. J Antimicrob Chemother 62:495–503.

Schaaper RM, Danforth BN, Glickman BW. 1986. Mechanisms of spontaneous mutagenesis: an analysis of the spectrum of spontaneous mutation in the Escherichia coli lacI gene. J Mol Biol 189:273–284.

Sreevatsan S, Pan X, Stockbauer KE, Williams DL, Kreiswirth BN, Musser JM. 1996. Characterization of rpsL and rrs mutations in streptomycin-resistant Mycobacterium tuberculosis isolates from diverse geographic localities. Antimicrob Agents Chemother 40:1024–1026.

Stover CK, Pham XQ, Erwin AL, Mizoguchi SD, Warrener P, Hickey MJ, Brinkman FS, Hufnagle WO, Kowalik DJ, Lagrou M, et al. 2000. Complete genome sequence of Pseudomonas aeruginosa PAO1, an opportunistic pathogen. Nature 406:959–964.

Trigos AS, Goudey BW, Bedo J, Conway TC, Faux NG, Wyres KL. 2021. Collateral Sensitivity to beta-Lactam Drugs in Drug-Resistant Tuberculosis Is Driven by the Transcriptional Wiring of BlaI Operon Genes. mSphere 6:e0024521.

Urbonavicius J, Qian Q, Durand JM, Hagervall TG, Bjork GR. 2001. Improvement of reading frame maintenance is a common function for several tRNA modifications. EMBO J 20:4863–4873.

Van Hofwegen DJ, Hovde CJ, Minnich SA. 2016. Rapid Evolution of Citrate Utilization by Escherichia coli by Direct Selection Requires citT and dctA. J Bacteriol 198:1022–1034.

Van TT, Minejima E, Chiu CA, Butler-Wu SM. 2019. Don’t Get Wound Up: Revised Fluoroquinolone Breakpoints for Enterobacteriaceae and Pseudomonas aeruginosa. J Clin Microbiol 57.

Wendrich TM, Blaha G, Wilson DN, Marahiel MA, Nierhaus KH. 2002. Dissection of the mechanism for the stringent factor RelA. Mol Cell 10:779–788.

